# Demixing of bacterial CipA and CipB proteins in mammalian cell cytosol provides an orthogonal self-assembly platform for producing isolable multi-phase intracellular crystals

**DOI:** 10.64898/2026.05.10.724141

**Authors:** Haruki Hasegawa, Songyu Wang, Emma Pelegri-O’Day

## Abstract

Crystalline inclusion proteins CipA and CipB from *Photorhabdus luminescens* serve as versatile scaffolds for clustering genetically fused heterologous enzymes into crystalline inclusion bodies. Although engineered Cip crystals are known to function as solid biocatalysts for improving metabolite production in bacterial cells, the phase separation behavior of Cip proteins in non-bacterial cellular environments, as well as their biochemical attributes in a soluble, non-crystalline state, remain poorly understood. This study demonstrates that CipA and CipB efficiently undergo crystallization in the cytosol of human embryonic kidney cells both at normal and hypothermic cell culture conditions. Within 72 hours post-transfection, CipA and CipB become the most abundant proteins in transfected cells and produce distinctive cytosolic crystals often exceeding 10 μm at least in one of the dimensions. Co-expression of CipA and CipB drives spontaneous demixing into two distinct crystal populations, and the orthogonality is maintained even when an unrelated third protein crystallizes in the same cytosol, permitting three crystal types to coexist simultaneously. Intracellular crystals are readily isolable from cells, and once purified, these crystals are stable under physiological pH conditions. However, CipA and CipB show notable differences in their crystal dissolution kinetics and protein oligomerization states when solubilized under acidic or alkaline conditions. These findings suggest that CipA/CipB forms a robust orthogonal self-assembly pair and establish CipA/CipB crystals as an efficient platform for producing biochemically programmable intracellular crystals. These properties should extend the Cip-based scaffolding approach to mammalian cell systems for synthetic biology applications.

## 1. Introduction

CipA and CipB are small hydrophobic crystalline inclusion proteins (Cip) expressed in the bioluminescent Gram-negative bacterium *Photorhabdus luminescens* [1]. *P. luminescens* forms a symbiotic relationship with the entomopathogenic nematode *Heterorhabditis bacteriophora* by residing within the nematode’s intestine and utilizing this association for infectious transmission [2, 3]. In return, *P. luminescens* supplies nutrients required for nematode reproduction and produces toxins and enzymes that mediate insect host killing [2, 3]. In *Photorhabdus* bacteria, expression of CipA and CipB protein is induced during exponential growth phase and peaks at the stationary phase when two distinct intra-cytoplasmic crystalline inclusions are produced. Rectangular Type I crystals are composed of CipB, while bipyramidal Type II crystals consist of CipA and, together, these two proteins can account for up to 40% of the total protein mass in bacteria [1, 4]. Insertional inactivation studies of the *cipA* or *cipB* genes showed these crystals do not act directly as insecticides, but function instead as nutrient reservoirs that support symbiotic nematode development [1, 3–5].

The CipA polypeptide is composed of 104 amino acids, is rich in methionine (14 residues), and contains four cysteine residues. CipB consists of 100 amino acids, has a high leucine content (18 residues), but lacks any cysteine residues. While CipA and CipB share roughly 25% of sequence identity (∼49% similarity) with each other, they have no sequence homology to any other known proteins, indicating that they constitute a unique protein class of their own [1]. The atomic structure of CipA was recently determined at a resolution of 2.11 (PDB ID: 7XHS), using nanocrystals generated by a cell-free translation system [6]. The CipA monomer consists of an N-terminal arm and a globular oligonucleotide/oligosaccharide-binding fold domain, which resembles the pentameric B subunit of heat-labile enterotoxin type IIB (PDB ID: 1QB5). Within crystals produced by the cell-free system, CipA monomers assemble through non-covalent interactions into tetramers with four-fold symmetry, which then serve as the basic units for CipA crystal formation [6]. These tetramer units are organized into multilayered structures and form a crystal lattice with porous spaces that can potentially accommodate guest macromolecules [6]. The monomeric structure and oligomeric state of the CipB protein remain uncharacterized to date.

Because of their ability to produce crystalline inclusions in bacteria, CipA and CipB crystals have drawn attention not only as novel biomaterials but also as versatile scaffolds capable of clustering and packaging foreign proteins into crystalline inclusions. In a pioneering work, Jung’s group [7] demonstrated that reporter proteins such as GFP (∼27 kDa) and LacZ (∼116 kDa) can produce optically refractile inclusions in *E. coli* when genetically fused to CipA and CipB sequences. The crystal-like inclusions can also be isolated from bacteria without losing the green fluorescence or β-galactosidase activity [7]. This proof-of-concept study prompted numerous follow-up investigations into Cip-based enzyme clustering and substrate-channeling approaches for engineering bacterial metabolic pathways. Notable examples include the clustering of five different enzymes for violacein biosynthesis [7], the use of exogenous enzymes for pyrogallol production [8], and the sequential conversion of valine to isobutyraldehyde through tandemly fused enzymes [9]. Clustering cytochrome P450 enzymes with partner reductases facilitated the production of pharmaceutical and nutraceutical compounds such as lutein, (+)-nootkatone, apigenin and L-DOPA [10]. Cip scaffolding also improved the synthesis of 3′-Phosphoadenosine-5′-phosphosulfate from ATP and sulfate [11], UDP-GlcNAc and UDP-GlcA [12], chiral alcohols [13], L-histidine [14], and Amaryllidaceae alkaloids [15]. The versatility of the Cip scaffold also enabled the formation of solid materials that can capture and recover rare earth elements [16] outside the biological context. When appropriately designed or engineered, CipA/CipB scaffolds therefore have the potential to produce a wide range of genetically encodable, isolable biocatalytic crystalline materials capable of fulfilling desired biochemical functions. To date, however, all successful applications have been limited to bacterial host cell systems.

This study demonstrates that CipA and CipB proteins, originating from *P. luminescens* and *P. laumondii*, can robustly produce cytosolic crystals in the human embryonic kidney cell line HEK293. Within 72 hours of transgene expression, CipA and CipB become the most abundant cellular proteins, and the crystal size frequently exceeds 10 μm at least in one of the dimensions, which is substantially larger than the dimensions of the bacterial cell. Upon co-expression, CipA and CipB undergo phase separation driven by strong homotypic interactions and produce two distinct crystal populations simultaneously. The orthogonality of CipA/CipB crystals is maintained even when an unrelated third protein undergoes crystallization within the same cytosolic compartment, allowing for the coexistence of three distinct crystal types simultaneously. Isolated intracellular crystals of CipA and CipB remain stable in aqueous environments at physiological pH. However, CipA and CipB show distinct properties regarding crystal dissolution kinetics and protein oligomerization status under acidic and alkaline conditions. Our findings reveal that the CipA/CipB pair constitutes a robust orthogonal self-assembly system and establish CipA/CipB crystals as an efficient platform for producing biochemically programmable intracellular crystals that can extend the Cip-scaffolding approach into mammalian cell systems.

## 2. Results

### 2.1. Overexpression of CipA and CipB from *Photorhabdus luminescens* leads to robust protein crystallization in the cytosol of mammalian cells

Because prokaryotic and eukaryotic cells differ in size, intracellular organization, and translational regulation, it remains unclear how these environments influence protein phase separation. Considering their recognized importance as crystalline scaffolds for enzyme immobilization, substrate channeling, metabolic engineering and crystalline biocatalyst production [7–15], we selected CipA and CipB from the entomopathogenic bacterium *Photorhabdus luminescens* as model proteins to investigate how their crystallization behaviors are maintained or modified within mammalian cellular environments.

To improve heterologous expression of bacterial proteins in mammalian cells, the nucleotide sequences encoding CipA and CipB were codon-optimized for human cell translation (see Materials and Methods). Within 72 hours post-transfection, recombinantly expressed CipA and CipB became the most abundant proteins in HEK293 cells (Fig. 1G, lanes 1 and 2, arrowhead near 10 kDa range). While native CipA forms bipyramidal and equiangular hexagonal crystals in stationary-phase *P. luminescens* cytoplasm [1, 4], the recombinant version of CipA in HEK293 cells produced a mixture of square, oblong hexagon, and four-pointed star-shaped crystals that measured ∼10 μm or larger in size by 3 days after transfection (Fig. 1, A-B). Similarly, while native CipB typically produces square and rectangular crystals in *P. luminescens* [1, 4], recombinant CipB generated regular hexagonal crystals in HEK293 cells (Fig. 1, D-E). In differential interference contrast (DIC) microscopy, these CipB crystals appeared as transparent hexagonal “empty space” in the mammalian cell cytoplasm. Similar crystal characteristics have been reported before for human NEU1 crystals that present as cubic “void” space in the endoplasmic reticulum (ER) lumen [17].

**Figure 1.**
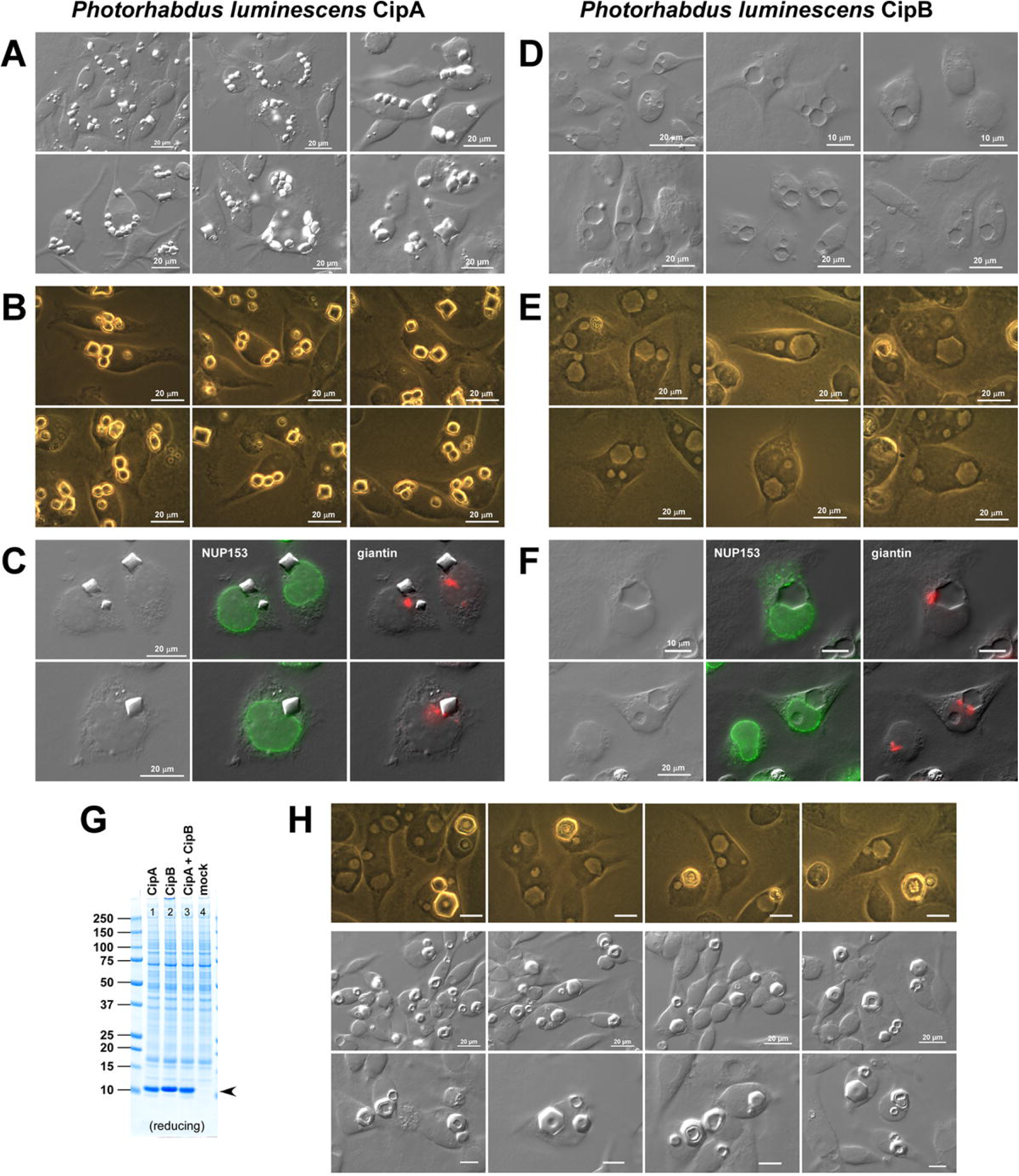
Recombinantly overexpressed *Photorhabdus luminescens* CipA and CipB undergo crystallization in the cytosol of mammalian cells. HEK293 cells overexpressing *P. luminescens* CipA (left panels, A-C) and CipB (right panels, D-F) were fixed and imaged 3 days after transfection. (A, D) Differential interference contrast (DIC) micrographs. (B, E) Phase contrast micrographs. (C, F) Immunofluorescent micrographs of transfected cells that were immunostained with mouse anti-NUP153 (shown in green) and rabbit anti-giantin (shown in red). (G) Coomassie blue-stained SDS-PAGE gel for whole cell lysates prepared from HEK293 cells expressing *P. luminescens* CipA (lane 1), CipB (lane 2), CipA and CipB (lane 3), and empty vector (lane 4). Cell lysates were prepared 72 hours after transfection. CipA and CipB proteins (∼11 kDa) are indicated by an arrowhead. (H, top row) Transfected cells imaged by phase contrast at day 4 reveal vacuolated donut-like CipB crystals. (H, second and third rows) DIC images at day 5 post-transfection where CipB crystal vacuolization is prevalent. Unlabeled scale bars represent 10 µm.

With extended cell culture beyond 4 days post-transfection, the initially intact hexagonal CipB crystals began to develop central vacuoles, peripheral cracks, and eventually showed severe deterioration, suggesting an inherent instability of the CipB crystalline lattice within cells (Fig. 1, H). Consistent with this observation, CipB crystals were previously found susceptible to pronase digestion, whereas CipA crystals were mostly protected [4].

Immunofluorescence staining using a nuclear membrane marker NUP153 and a Golgi marker giantin revealed that both CipA and CipB crystals were located in the cytosol of HEK293 cells, and excluded from the nucleus and Golgi apparatus, as expected from the absence of identifiable organelle targeting signals (Fig. 1, C, F). Notably, these CipA and CipB crystals were significantly larger than the crystals present in the cytoplasm of *P. luminescens* bacterium [1, 4]. In fact, the size of the crystals in HEK293 cells exceeded the dimensions of the bacterium itself, which measures 0.5–1.0 μm in diameter and 2.0–5.0 μm in length. Despite being composed of identical polypeptides in both human HEK293 and *P. luminescens* cells, the higher biosynthetic capacity and larger size of mammalian cells may have enabled CipA/CipB crystals to grow well beyond the spatial constraint of their native bacterial environment.

### 2.2. Two morphologically different protein crystals are simultaneously maintained in HEK293 cells upon CipA and CipB co-expression

An expanding number of proteins are known to crystallize within cells [18, 19], and research shows that as many as four distinct crystals can be recombinantly induced simultaneously inside one mammalian cell [17, 20], or even within a single organelle [21]. Of all reported cases of intracellular crystallization across organisms and experimental systems, the CipA/CipB protein pair from *P. luminescens* represents a unique case in which two distinct, endogenously expressed proteins produce crystals and naturally coexist in the bacterial cytoplasm [1, 4].

To examine whether CipA and CipB can spontaneously demix and stably coexist as independent crystal populations in the cytosol of mammalian cells (thereby phenocopying their behavior in bacterial cells), both genes were co-transfected into HEK293 cells at a 1:1 DNA ratio. Although this approach reduced the gene dosage of each construct by 50% (as the total DNA amount was kept constant), transfected HEK293 cells achieved high protein expression levels, resulting in CipA and CipB being the most abundant proteins in the cell (Fig. 1G, lane 3). Under these co-expression conditions, the transfected HEK293 cells started harboring two different types of protein crystals simultaneously, one made of CipA and the other of CipB. These crystals coexisted stably while each crystal type retained its characteristic morphology (Fig. 2, A, B). As observed during individual expression, both crystal types were excluded from the nucleus and located in the cytosolic compartment (Fig. 2C). With only 26.2% sequence identity and 46.6% homology between CipA and CipB (Suppl. 1A), these two proteins may not interact, if at all, with each other in the native cytoplasm of *P. luminescens* and in the heterologous environment of the HEK293 cell cytosol. In other words, homotypic interactions among CipA or CipB appeared to outweigh heterotypic and nonspecific interactions, thereby driving spontaneous phase separation into distinct crystalline inclusion bodies.

**Figure 2.**
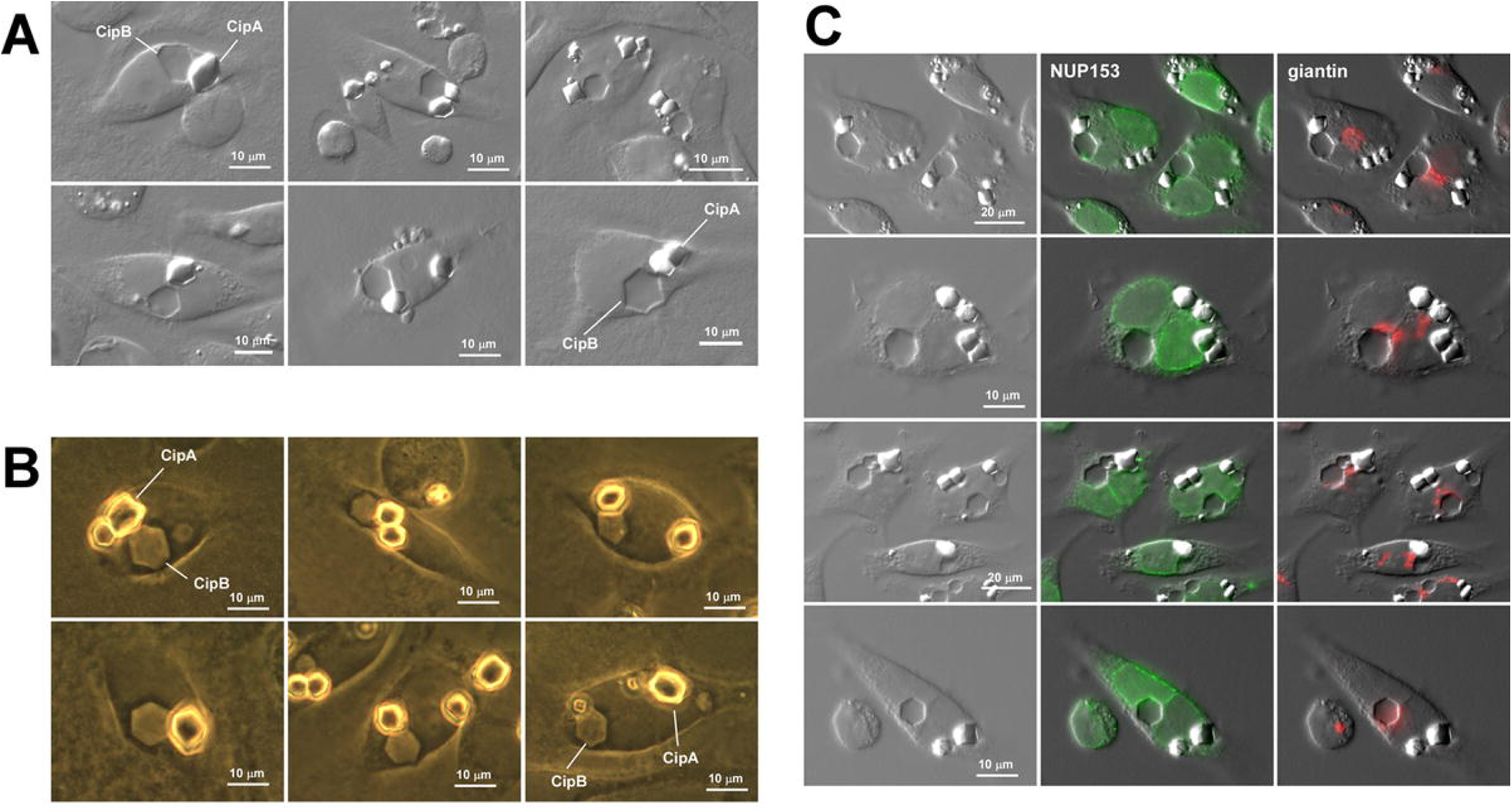
Simultaneous formation of two distinct crystal types in the cytoplasm of HEK293 cells co-expressing CipA and CipB. (A, B) DIC and phase contrast images of HEK293 cells that simultaneously house CipA and CipB protein crystals on day 4 post-transfection. CipA and CipB crystals are labeled in representative images. (C) Immunofluorescent micrographs of co-transfected cells fixed on day 4 post-transfection. The cells were permeabilized and immunostained with mouse anti-NUP153 (shown in green) and rabbit anti-giantin (shown in red).

### 2.3. CipA and CipB from the closely related bacterium *Photorhabdus laumondii* also undergo crystallization in the cytosol of mammalian cells

According to the UniProt database, there is another listed pair of CipA and CipB proteins from a different bacterial species within the same *Photorhabdus* genus, namely, *Photorhabdus laumondii* [22]. CipA (UniProt Q7N6H4) and CipB (UniProt Q7N9Z4) from *P. laumondii* share 92.3% and 95.0% sequence identity with their *P. luminescens* counterparts, respectively (see Suppl. 1, C-D). As observed in *P. luminescens*, the sequence similarity between CipA and CipB within *P. laumondii* remains low, with only 24.0% identity and 46.2% homology (Suppl. 1B).

When *P. laumondii* CipA was overexpressed alone, the intracellular CipA crystal population showed square, bipyramidal, or four-pointed star-like contours (Fig. 3A, top two rows; Fig. 3E, lane1) and they were localized in the cytosol (Fig. 3A, third and fourth rows). *P. laumondii* CipB produced hexagonal protein crystals; however, these crystals showed irregular shapes and frequent central vacuolation as early as 3 days post-transfection (Fig. 3B, first and second rows; Fig. 3E, lane 2). The extent of vacuolization varied considerably, and occasionally, these vacuolated inclusions were found in the nucleoplasm in addition to the cytosol (Fig. 3B, third and fourth rows). Compared to *P. luminescens* CipB crystals, those produced by *P. laumondii* deteriorated more rapidly, possibly reflecting even greater packing instability or lower tolerance to packing defects.

**Figure 3.**
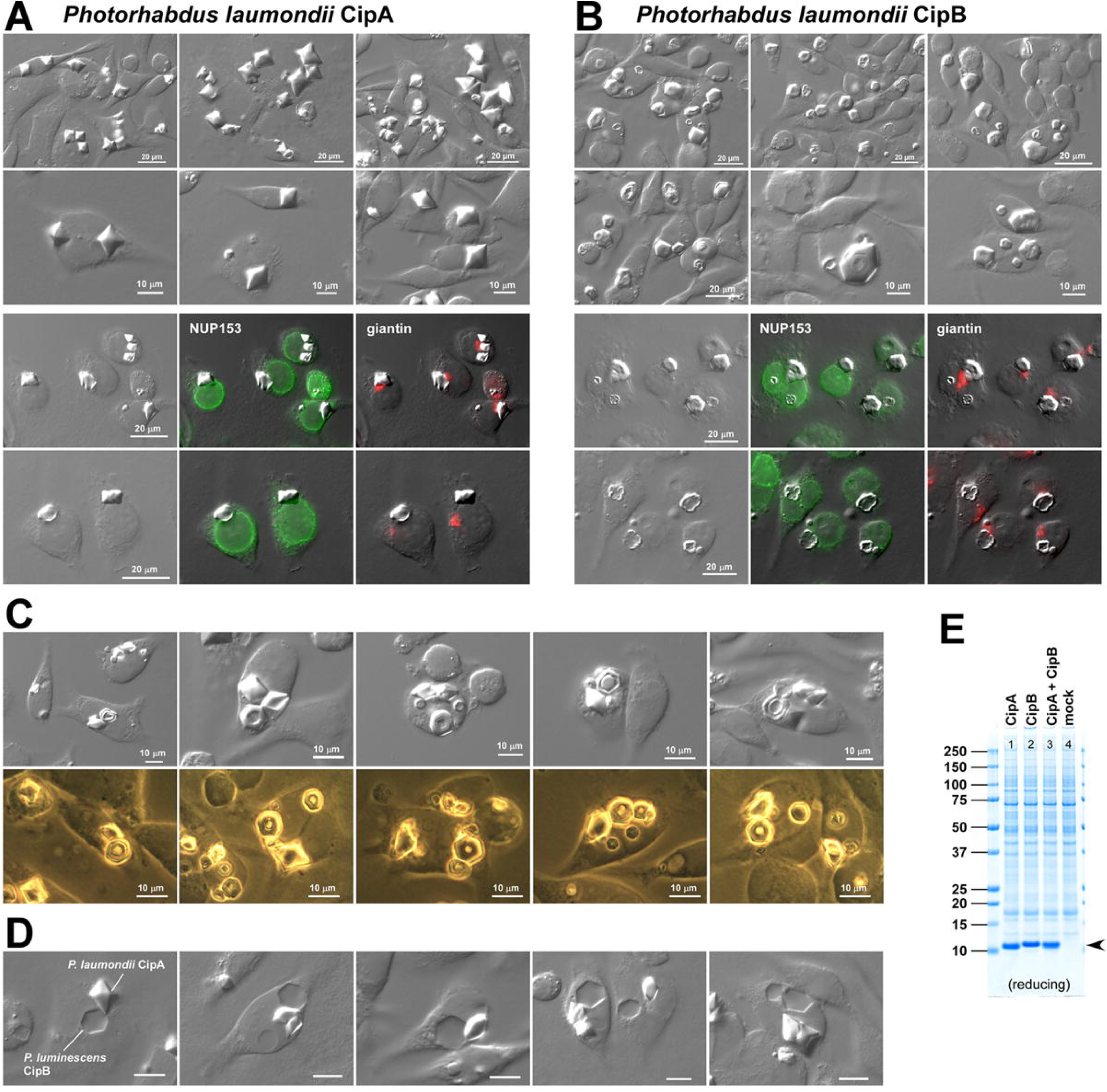
Co-expression of *Photorhabdus laumondii* CipA and CipB leads to simultaneous formation of two distinct crystal inclusion bodies in mammalian cell cytoplasm. (A, B) DIC images (first and second rows) and immunofluorescent images (third and fourth rows) of transfected HEK293 cells expressing *P. laumondii* CipA (left panels) and CipB (right panels). On day 4 after transfection, cells were fixed and immunostained using mouse anti-NUP153 (green) and rabbit anti-giantin (red). (C) DIC images (top row) and phase contrast images (second row) of HEK293 cells that simultaneously house CipA and CipB protein crystals. Cells were fixed and imaged on day 4 post-transfection. (D) DIC micrographs of HEK293 cells co-expressing *P. laumondii* CipA and *P. luminescens* CipB. Cells were fixed and imaged on day 4 post-transfection. Unlabeled scale bars represent 10 µm. (E) Coomassie blue-stained SDS-PAGE gel for whole cell lysates prepared from HEK293 cells expressing *P. laumondii* CipA (lane 1), CipB (lane 2), CipA and CipB (lane 3), and empty vector (lane 4). Cell lysates were prepared at 72 hours post-transfection. CipA and CipB proteins (∼11 kDa) are indicated by an arrowhead.

Although *P. laumondii* CipB crystals exhibit instability, both CipA and CipB proteins showed robust expression levels whether transfected individually or co-transfected at a 1:1 DNA ratio (Fig. 3E, lanes 1-3). The high protein levels at steady state indicate that cellular protein turnover mechanisms are likely not significantly contributing to the CipB crystal quality deterioration. Additionally, when co-expressed, *P. laumondii* CipA crystals and the unstable CipB crystals could coexist in the same cytosolic space without affecting each other (Fig. 3C). Likewise, co-expression of *P. laumondii* CipA with *P. luminescens* CipB, despite originating from different bacterial species, resulted in the stable formation of two distinct crystal types in the cytosol (Fig. 3D), suggesting that homotypic protein interactions predominate over other associative forces.

### 2.4. Intracellular CipA and CipB protein crystals retain their characteristic morphology even under hypothermic conditions

In their natural habitats, both *P. luminescens* and its symbiont nematode *Heterorhabditis bacteriophora*, as well as their insect prey complete their life cycles under ambient temperature conditions [2, 3]. To determine if there are any temperature-dependent phase behaviors or temperature-gated folding steps critical for crystallization, CipA– and CipB-transfected HEK293 cells were transferred to a 27 °C CO_2_ incubator 24 hours after transfection (before the crystals begin to appear). The growth of intracellular crystals was monitored for 5 days following the temperature shift.

CipA and CipB from *P. luminescens* retained crystallizing propensity at 27 °C, with the resulting crystals preserving their characteristic morphology in both single gene expression and co-transfection experiments (Fig. 4A). However, crystal growth appeared slower at 27 °C, as indicated by smaller crystal sizes, which was most likely caused by reduced metabolic activity and lower rates of protein synthesis at this hypothermic temperature. Comparable results were observed for CipA and CipB from *P. laumondii* and the rapid crystal deterioration of *P. laumondii* CipB was not prevented by this temperature setting (Fig. 4B). Unlike firefly luciferase, which crystallizes only at ambient temperature rather than at mammalian body temperature [23], the characteristic phase separation behavior of CipA and CipB was not restricted to a narrow temperature range.

**Figure 4.**
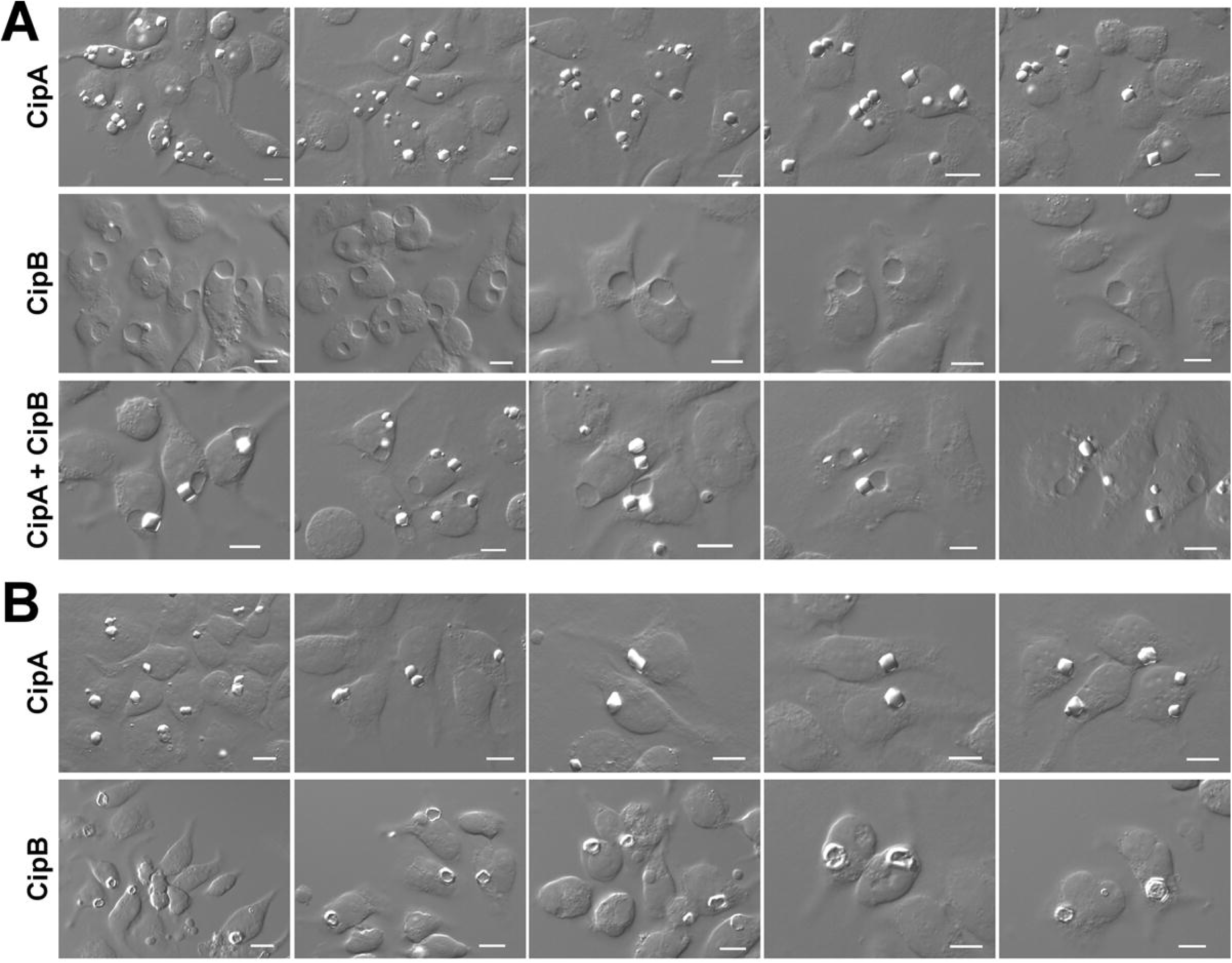
CipA and CipB proteins crystallize in HEK293 cells even under hypothermic cell culture conditions. (A) DIC micrographs of HEK293 cells expressing *P. luminescens* CipA (top row), CipB (second row), and CipA and CipB co-transfection (third row). At 24 hours post-transfection, cells were seeded onto glass coverslips and transferred to an incubator maintained at 27 °C, 5 % CO_2_. After incubating 72 hours at the hypothermic cell culture conditions (namely, 96 hours post-transfection), cells were fixed and imaged. (B) HEK293 cells expressing *P. laumondii* CipA (top row) and CipB (second row) were similarly fixed and imaged after 72 hours of hypothermic cell culture. Unlabeled scale bars represent 10 µm.

### 2.5. CipA and CipB crystals remain stable and independent even in the presence of unrelated proteins simultaneously undergoing crystallization

To evaluate the robustness of the CipA and CipB intracellular crystallization process, we tested whether co-expressing another cytosolic phase-separating protein would disrupt their crystal formation, directly or indirectly. The rationale is that if self-association of CipA or CipB is labile and can be disrupted by the newly-introduced second protein, this new interaction may lead to changes in crystal morphology or potentially inhibit crystal formation entirely.

In the initial set of evaluations, we examined how the crystallization of *P. luminescens* CipA and CipB changes when a highly homologous protein is present in the shared cytosolic environment. Specifically, *P. laumondii* CipA was co-transfected into cells expressing *P. luminescens* CipA. Given their 92.3% sequence identity, these proteins may substitute for each other during crystal assembly. The co-expression of two related CipA proteins resulted in abundant intracellular crystals (Fig. 5A). However, due to their intrinsically similar crystal morphologies, it was not possible to distinguish homotypic crystals comprised solely of one CipA variant from potential heterotypic co-crystals (Fig. 5A). This limitation also precluded assessment of any dominance between the two CipA variants. Comparable challenges arose with cells co-expressing *P. luminescens* CipB and *P. laumondii* CipB, which share 95.0% sequence identity (Fig. 5B). Although this outcome was not unexpected, introducing a closely related crystallizing protein made it impossible to determine whether, or how, the crystallization of the first protein was influenced. This was due to limitations of our label-free detection methods, which cannot reveal the distribution of individual protein molecules within the crystals.

**Figure 5.**
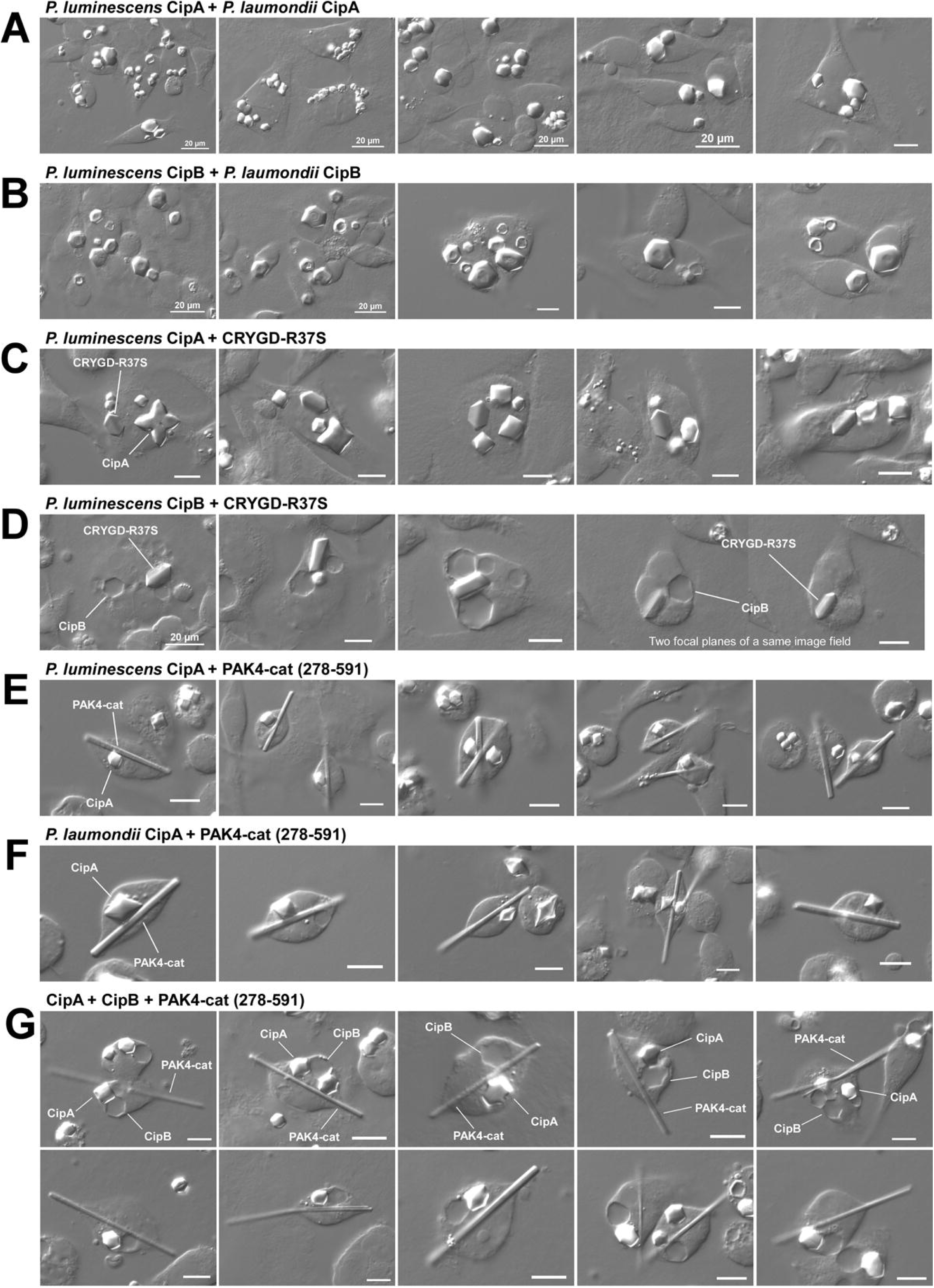
CipA and CipB crystals remain stable and independent in the presence of unrelated proteins that also produce crystals in the cytosol. DIC micrographs of HEK293 cells fixed and imaged on day 4 post-transfection. Cells were transfected to co-express the following protein combinations: (A) *P. luminescens* CipA and *P. laumondii* CipA, (B) *P. luminescens* CipB and *P. laumondii* CipB, (C) *P. luminescens* CipA and CRYGD-R37S, (D) *P. luminescens* CipB and CRYGD-R37S, (E) *P. luminescens* CipA and PAK4-cat (278-591), (F) *P. laumondii* CipA and PAK4-cat (278-591), (G) *P. luminescens* CipA, *P. luminescens* CipB, and PAK4-cat (278-591). Representative protein crystals are labeled with protein names. Unlabeled scale bars represent 10 µm.

To circumvent observed assay limitations, we repeated the experiments using the model proteins that are biologically and biochemically unrelated to CipA or CipB. A cataract-associated mutant of human γ-crystallin D carrying the R37S point mutation (CRYGD-R37S) was shown previously to form distinctive hexagonal crystals in the cytosol [17]. Thanks to its sequence dissimilarity with *P. luminescens* CipA and CipB, co-expression of CRYGD-R37S with CipA or CipB resulted in the formation of two morphologically different crystal populations in the cytosol, with no apparent interference between them (Fig. 5, C-D). Similarly, co-expression of CipA (*P. luminescens*) or CipA (*P. laumondii*) with PAK4-cat (278-591), another unrelated protein known to generate rod-shaped cytosolic crystals and serve as an enzyme scaffold [24], yielded two independent crystal populations, each retaining its distinct morphological identity without affecting one another (Fig. 5, E-F). Furthermore, simultaneous co-expression of three proteins, i.e., CipA and CipB from *P. luminescens* and PAK4-cat (278-591), led to the formation of three discrete phase-separated crystal populations within the same cytosolic environment, with no evidence of mutual inhibition or interference during crystallogenesis (Fig. 5G). Collectively, these results demonstrate that CipA and CipB crystals maintain stability and individuality even in the presence of unrelated crystallizing proteins, highlighting their strong self-association properties in the crowded environment of mammalian cell cytosol.

### 2.6. CipA/CipB crystals remain stable at physiological pH, but show distinct dissolution profiles in acidic or basic environments

To biochemically characterize CipA and CipB proteins, intracellular protein crystals were extracted from crystal-housing HEK293 cells through detergent cell lysis and subsequent crystal pellet washing (see Materials and Methods). This simple method yielded highly purified CipA and CipB crystals for both *P. luminescens* and *P. laumondii* species (Fig. 6, A-B). As anticipated, the morphology of isolated protein crystals matched their original intracellular crystal forms.

**Figure 6.**
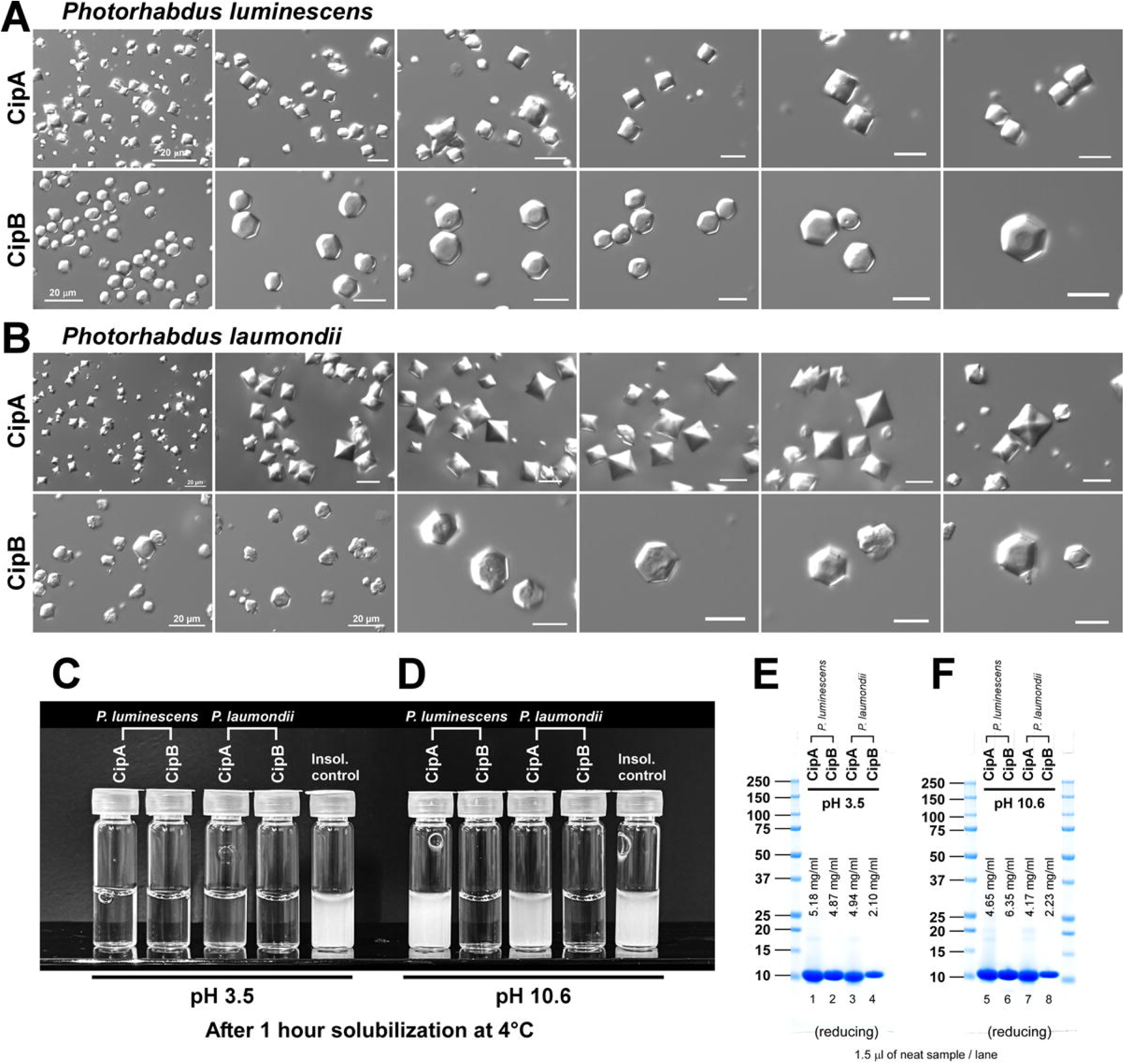
Isolation and solubilization of intracellular CipA and CipB crystals. (A, B) DIC micrographs of isolated intracellular crystals at different scales. CipA and CipB crystals shown in panels A and B are the proteins from *P. luminescens* and *P. laumondii*, respectively. Unlabeled scale bars represent 10 µm. (C, D) Solubility of CipA and CipB crystals at different pH settings. Purified intracellular crystals of *P. luminescens* and *P. laumondii* CipA/CipB obtained above were resuspended in pH 3.5 buffer (panel C) or in pH 10.6 buffer (panel D). After static storage at 4 °C in the dark for 1 hour, the solubilized protein samples were thoroughly mixed. A portion was then transferred to glass vials to be photographed. (E, F) After 7 days of static storage at 4 °C in the dark with daily gentle pipetting and mixing, fully dissolved CipA and CipB proteins at pH 3.5 (panel E) and pH 10.6 (panel F) were analyzed by SDS-PAGE under reducing conditions. Protein concentration obtained by OD_280_ measurement is shown in each lane. The presented data are representative of results obtained from three independent experiments.

The isolated crystals remained stable and resisted solubilization in Hepes-buffered saline (HBS) at physiological pH for at least up to 19 days at 4 °C. In contrast, all four crystal types dissolved easily in an acidic buffer (pH 3.5) within 1 hour at 4 °C (Fig. 6C). In a basic pH buffer (pH 10.6), only the CipB crystals dissolved within 1 hour, while CipA crystals remained insoluble during the same period (Fig. 6D). CipA crystals required an extended incubation period of ∼3–4 days to become soluble in the pH 10.6 buffer. To assess the quality and quantity of dissolved CipA and CipB proteins, SDS-PAGE was performed after a 7-day solubilization process in the respective pH settings. Under reducing conditions, solubilized crystals resolved into a single major band, irrespective of the solubilization pH (Fig. 6 E-F).

Mass spectrometry analysis under reducing conditions revealed an increase of 32 Daltons and 16 Daltons in protein mass for *P. luminescens* CipA and CipB, respectively. Likewise, for *P. laumondii* proteins, an increase of 32 Daltons was observed for CipA and 16 Daltons for CipB. These mass shifts suggest that one or two amino acid residues have undergone oxidation or hydroxylation per molecule. No additional chemical or post-translational modifications were identified. The crystal isolation and solubilization protocol (normalized by starting cell number and final solubilization volume) consistently yielded protein preparations at a concentration of ∼5 mg/ml (1 ml each), with the exception of *P. laumondii* CipB crystals, which resulted in yields roughly 50% lower than the other three (see Fig. 6, E-F). Based on these values, and assuming 100% of cells housed protein crystals at 72 hours post-transfection and that there was no yield loss during crystal isolation and solubilization, the obtained yield (∼5 mg/ml, 1 ml each) equates to approximately 25 pg of CipA or CipB crystals per cell. However, in practice, only about 90% of the total cell population contained intracellular crystals of varying sizes and numbers, and some degree of protein loss is expected; the true value is likely higher than 25 pg crystals/cell. Nonetheless, three conclusions can be drawn: (a) pure intracellular CipA and CipB crystals are readily isolable from transfected cells, (b) isolated crystals can be stored stably in a physiological pH buffer for an extended period, and (c) the crystals are solubilized by subjecting them to either acidic or basic pH conditions as needed.

### 2.7. CipA protein undergoes gradual intermolecular disulfide bond formation upon solubilization in basic pH buffer

Solubilization of CipA and CipB crystals at two different pH conditions yielded highly pure protein preparations (Fig. 6 E-F). Here, the same protein samples were re-examined using SDS-PAGE under non-reducing conditions. This analysis revealed that CipA and CipB have different susceptibilities to air oxidation depending on the pH settings. Both CipA and CipB remained as monomers after 7 days at pH 3.5 (Fig. 7A, lanes 1-4), and CipB also remained monomeric at pH 10.6 (Fig. 7A, lanes 6 and 8). However, CipA formed covalently linked dimers and trimers via disulfide bonds only at pH 10.6 (Fig. 7A, lanes 5 and 7, red arrowheads). Notably, this adduct formation was absent in CipB, likely due to the lack of cysteine residues in its primary sequence, unlike *P. luminescens* CipA and *P. laumondii* CipA that contain four and five cysteine residues, respectively (refer to Suppl. 1C).

**Figure 7.**
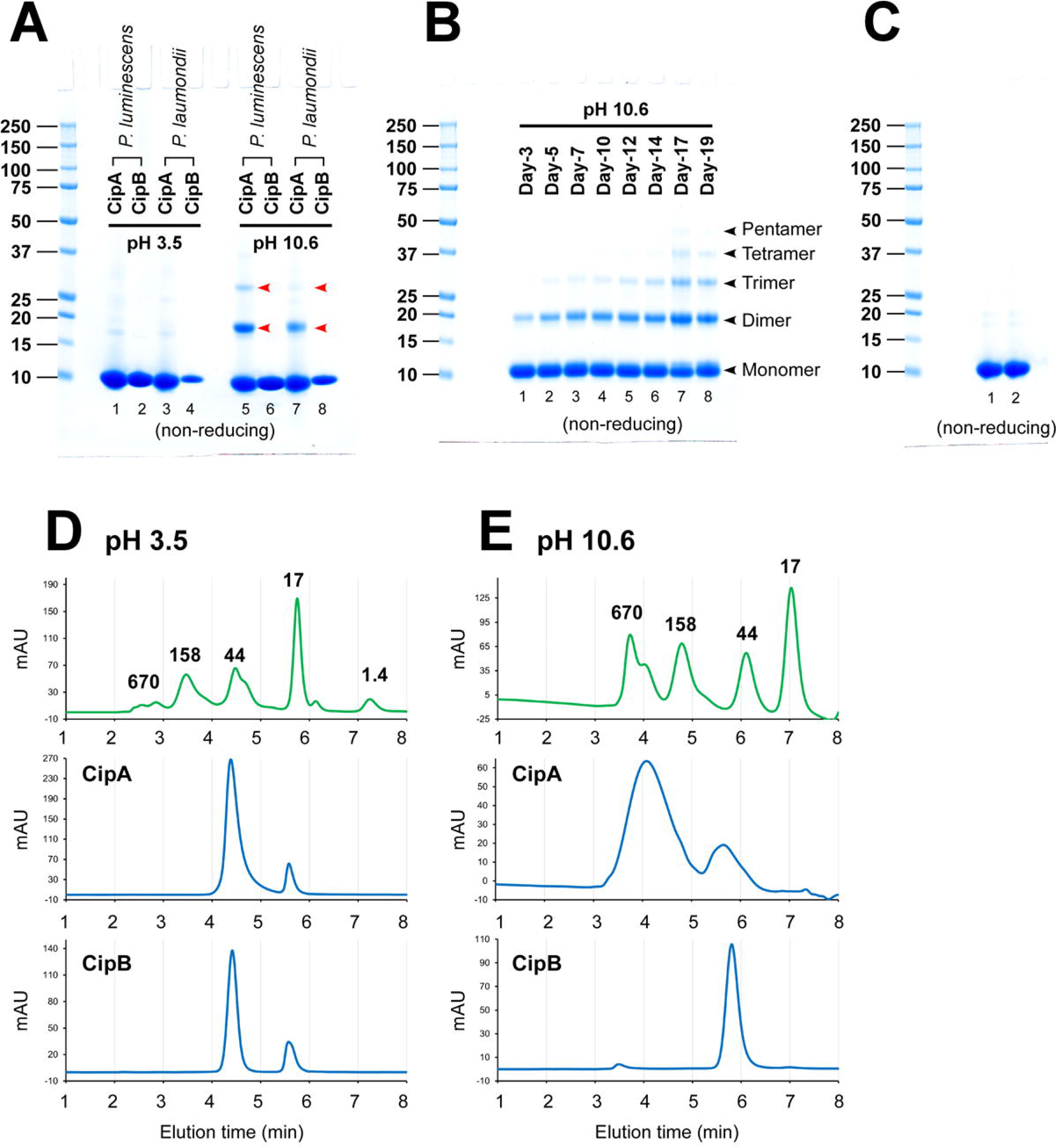
Oligomeric states of solubilized CipA and CipB proteins are strongly influenced by the solution pH. (A) After 7 days of solubilization at 4 °C in the dark using pH 3.5 buffer (lanes 1-4) and pH 10.6 buffer (lanes 5-8), fully dissolved CipA and CipB proteins were analyzed by SDS-PAGE under non-reducing conditions. Disulfide-linked dimer and trimer species found in CipA at pH 10.6 are indicated by red arrowheads (see lanes 5 and 7). (B) Time course of *P. luminescens* CipA covalent multimer formation at pH 10.6. From day 3 to day 19, an aliquot of CipA protein solubilized in pH 10.6 buffer was analyzed by SDS-PAGE under non-reducing conditions. Sample loading, 10 μg/lane. (C) *P. luminescens* CipA crystals stored in pH 7.4 buffer for 19 days were directly solubilized in non-reducing SDS sample buffer and analyzed by SDS-PAGE under non-reducing conditions. Lanes 1 and 2 are duplicated loading of the same sample. Loading 10 μg/lane. (D, E) Solubilized CipA and CipB proteins were analyzed under non-denaturing conditions by SEC. CipA and CipB crystals were solubilized in (D) pH 3.5 buffer and (E) pH 10.6 buffer and subjected to SEC using respective solubilization buffers at a flow rate of 0.45 ml/min. Protein loading was 10 μl each at ∼5 mg/ml concentration. The elution profile was monitored by UV light at 215 nm. AU, arbitrary unit.

To track the timing and the extent of cysteine oxidation during CipA crystal solubilization in pH 10.6 buffer, *P. luminescens* CipA protein quality was monitored by non-reducing SDS-PAGE up to day 19. As before, CipA crystals mostly dissolved after 3 days, with some covalent dimers already formed (Fig. 7B, lane 1). From day 5 onwards, dimer species increased steadily, and trimers began to appear (Fig. 7B, lanes 2-8). By day 17, faint bands of tetramers and pentamers were detectable (Fig. 7B, lanes 7 and 8), indicating slow progression of covalently linked multimer formation. The gradual nature of multimer formation suggests that the process is not enzymatically catalyzed but instead results from slow chemical oxidation mediated by dissolved oxygen in the aqueous buffer.

To determine if covalently adducted CipA species preexisted in the CipA crystals, the crystals were stored in HBS buffer (pH 7.4) for 19 days under identical storage conditions as above, thoroughly washed, then dissolved directly in non-reducing SDS sample buffer for SDS-PAGE analysis. No covalently linked CipA species were detected when directly dissolved from the crystals (Fig. 7C, lanes 1-2), suggesting that inter-molecular disulfide formation is promoted only after CipA is solubilized in pH 10.6 buffer. While the biological significance of CipA covalent multimerization at basic pH is unclear, identifying involved cysteine residues and managing the oxidation reaction will be important for protein scaffold engineering and product quality control.

### 2.8. Solubilized *P. luminescens* CipA and CipB proteins produce different types of non-covalently associated multimers under different pH environments

To further characterize the soluble forms of recombinant CipA and CipB proteins, the quality of CipA and CipB proteins was examined under non-denaturing conditions by size exclusion chromatography (SEC). At pH 3.5, both CipA and CipB behaved similarly and eluted as a major peak near the 44 kDa molecular weight marker (Fig. 7D), thereby indicating that these proteins form multimers at this pH. Likewise, the elution profiles revealed that CipA and CipB are present in oligomeric or aggregated forms under solubilization at pH 10.6. CipB was observed as a single, well-defined peak near the 44 kDa marker (Fig. 7E, bottom), whereas CipA resolved into two broad peaks; a minor peak eluted earlier than the 44 kDa sizing marker and a major broad peak found in the high molecular weight range (Fig. 7E, middle). Because the day 7 CipA protein sample used for this SEC analysis contained only a small fraction of covalently linked dimers and trimers (Fig. 7A, lane 5; 7B, lane 3), the results suggest that the majority of soluble CipA had already formed non-covalently associated higher-order aggregates in alkaline solutions before the covalent adducts gradually formed.

Since gel filtration profiles do not always reveal a protein’s exact oligomeric state, SEC-MALS analysis was performed to determine the molecular mass of the oligomeric CipA and CipB at pH 3.5 and pH 10.6. Under pH 3.5 solubilization conditions, the main CipA peak corresponded to a species of ∼39.8 kDa, likely representing a mixture of trimers and pentamers (Fig. 8A, Table-1). In contrast, the main peak for CipB was comprised of a ∼42.9 kDa species, most probably corresponding to tetramers (Fig. 8C, Table-1). When subjected to pH 10.6 solubilization, the prominent high molecular weight peak of CipA was determined as ∼117 kDa species, consistent with the decamer size, while the secondary peak was ∼25 kDa, indicative of dimeric species (Fig. 8B, Table-1). Under these same alkaline conditions, CipB remained as a ∼43.9 kDa tetrameric species (Fig. 8D, Table-1).

**Figure 8.**
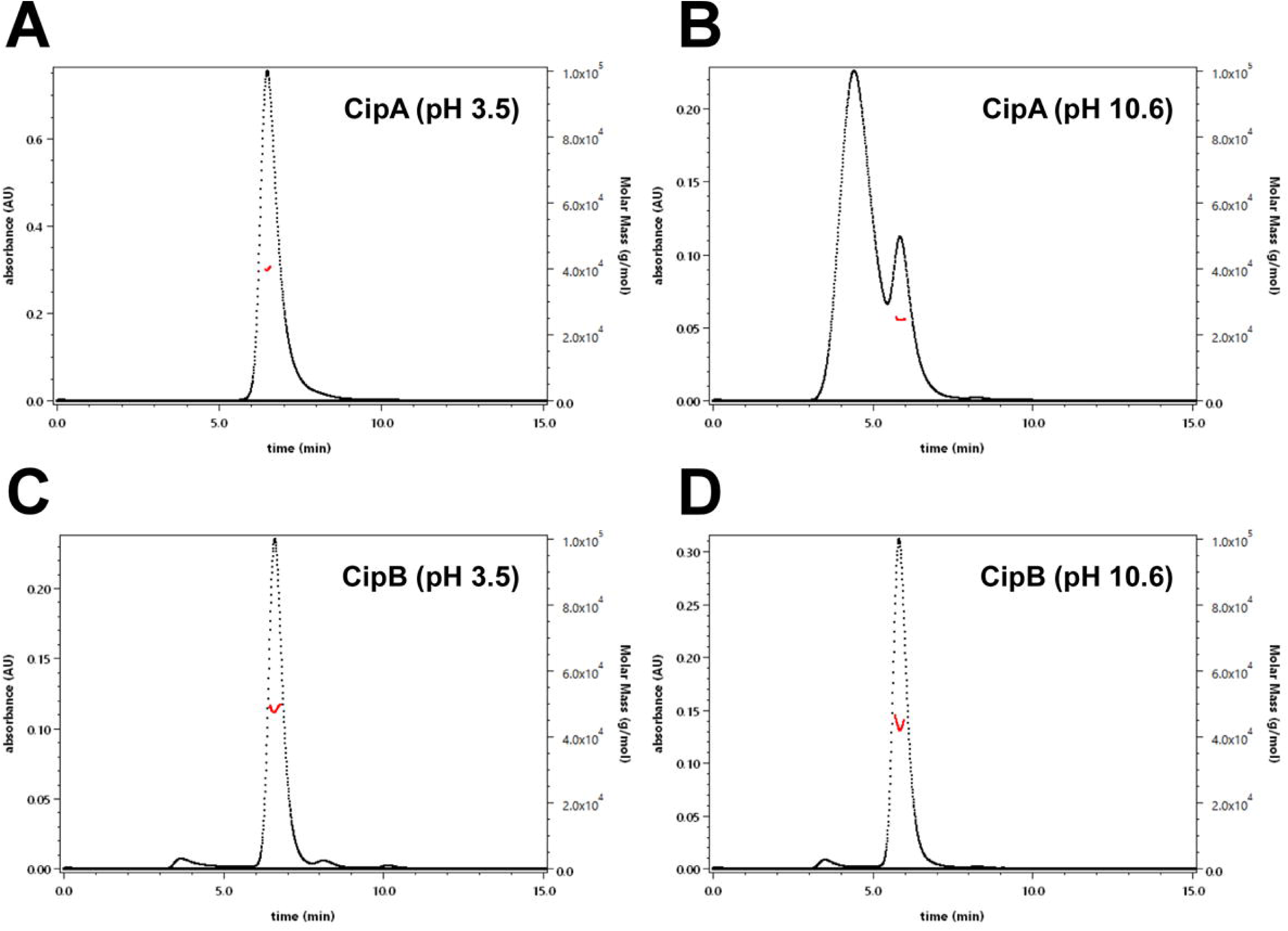
Molecular mass assignment for oligomeric CipA and CipB. Molecular mass of solubilized CipA and CipB was determined by SEC-MALS. Light scattering (left axis) and molecular mass (right axis) are plotted by black dots and red dots in each panel. (A) CipA at pH 3.5. (B) CipA at pH 10.6. (C) CipB at pH 3.5. (D) CipB at pH 10.6.

**Table-1.** Summary of SEC-MALS results for solubilized CipA and CipB proteins. The table presents the sequence-based calculated molecular masses, observed molecular masses, and proposed oligomeric states for CipA and CipB under pH 3.5 and pH 10.6 solubilization conditions.

|  |  | Calculated mass | Observed mass | Oligomeric state |
| --- | --- | --- | --- | --- |
| **pH 3.5** | CipA | 11.697 kDa | 39.80 kDa ± 8.4% | Trimer/Tetramer |
| | CipB | 11.315 kDa | 42.97 kDa ± 8.1% | Tetramer |
| **pH 10.6** | CipA [Peak 1] | 11.697 kDa | 117.50 kDa ± 10.0% | Decamer |
| | CipA [Peak 2] | 11.697 kDa | 24.98 kDa ± 8.5% | Dimer |
| | CipB | 11.315 kDa | 43.94 kDa ± 7.9% | Tetramer |

Although previous studies identified non-covalently assembled CipA tetramers as the basic building block of CipA crystals [6], our findings indicate that, upon solubilization under acidic or alkaline conditions, CipA proteins are capable of reorganizing into various oligomeric forms. Conversely, while the oligomeric state of CipB in its crystalline state remains unresolved, solubilized CipB consistently retains its tetrameric configuration at acidic and alkaline pH environments.

## 3. Discussion

### 3.1. CipA/CipB crystallization is driven by robust homotypic interactions in mammalian cells

Our study demonstrated that the bacterial crystalline inclusion proteins, CipA and CipB, retained a strong intrinsic propensity to self-associate and crystallize even when expressed in a eukaryotic cellular context using the human kidney cell line, HEK293. Both CipA and CipB accumulated to become the most abundant proteins in transfected HEK293 cells and resulted in the formation of readily identifiable crystals within the cytosolic compartment. Whether the substantial stockpiling of CipA/CipB is a consequence of the stabilizing/storage effects of protein crystallization, or whether sustained high-level synthesis is required to initiate crystallization, remains unclear. At present, this relationship presents a chicken-and-egg problem. The observed cytosolic localization in the eukaryotic environment was consistent with the absence of recognizable organelle-targeting signals such as an ER targeting signal peptide, a peroxisome targeting SKL motif, or a membrane anchor domain.

A particularly striking observation was the scale of crystal growth (with crystals reaching sizes greater than 10 μm within 72–96 hours after transgene expression) that well exceeded the dimensions of the bacterial cells. A plausible interpretation is that the substantially greater cell volume and higher biosynthetic capacity of mammalian cells mitigated geometric and resource limitations on crystal size, thereby permitting the continued deposition of proteins onto the expanding crystalline lattice beyond the physiological constraints imposed in bacterial cells. Notably, cell size is a key determinant of intracellular crystal size even among different mammalian cell types [17, 25]. Although high crystallization penetrance (∼90% of cells) and estimated crystal mass of ∼25 pg/cell were obtained in the HEK293 transient expression platform, even greater total crystal yield may be achieved by the stable expression of CipA and CipB in Chinese Hamster Ovary cells that are characterized by larger cell size and higher biosynthetic capacity [25].

Importantly, CipA and CipB maintained their crystallization propensity and characteristic crystal morphologies even when HEK293 cells were cultured at a hypothermic temperature of 27 °C. The apparent slow crystal growth under these conditions was attributed to reduced protein synthesis and lower metabolic activity at the lower temperature. The relative insensitivity of CipA/CipB crystallization to temperature, spanning a 10 °C range from 27 °C to 37 °C, suggests that their crystallization is not tightly regulated by temperature-sensitive folding kinetics or steps [23]. This permissive capacity enables CipA/CipB crystallization to be implemented in various recombinant host systems and under different cell culture conditions.

### 3.2. Purified CipA/CipB crystals maintain stability at physiological pH, while exhibiting distinct solubilization profiles under acidic and alkaline conditions

A practically important finding is that CipA and CipB crystals can be efficiently purified from mammalian cells by simple detergent cell lysis and low-speed centrifugation steps. The purified crystals retained their original morphology and remained stable outside the cellular environment when formulated in a physiological pH buffer, allowing the possibility of long-term crystal storage.

When subjected to acidic conditions (pH 3.5), both CipA and CipB crystals dissolved easily. However, CipA crystals exhibited markedly slower dissolution kinetics under basic conditions (pH 10.6), requiring ∼3–4 days for complete solubilization. This differential CipA dissolution behavior at alkaline pH is notable because a prior study using a bacterial expression system reported that isolated CipA and CipB crystal-like insoluble materials could be solubilized rapidly at both pH 4.0 and pH 11.0 [4]. The slower alkaline dissolution of CipA crystals produced in HEK293 cells may reflect differences in crystal packing, post-translational modification, or unique physicochemical attributes conferred by the mammalian cellular environment. Mass spectrometry indicated that, while a minor fraction of the native CipA produced in *P. luminescens* was acetylated at the N-terminus [4], CipA and CipB produced in HEK293 cells were characterized primarily by oxidation at one or two amino acid residues per molecule as their only detectable chemical modification. Crystallographic characterization of the crystalline lattice would clarify whether crystals formed in mammalian cells share lattice parameters with those produced in bacteria. The ability to selectively dissolve CipB crystals under alkaline conditions while retaining CipA crystals (at least transiently for a few days) can also be exploited as a biochemically programmable, controlled-release mechanism for cargo proteins genetically fused to Cip-based crystalline materials.

### 3.3. CipA forms non-covalent higher-order structures and time-dependent disulfide adducts at basic pH

When solubilized and examined under acidic conditions (pH 3.5), CipA and CipB were found to form trimers (with likely coexistence of tetramers) and tetramers, respectively. The observation of trimeric soluble CipA protein species was unexpected considering that Ueno’s X-ray crystallography data identified tetramers as the building units of CipA crystals generated via a cell-free protein synthesis platform [6]. This suggests that CipA may exhibit variable oligomeric valency, adopting different forms depending on whether it is in a soluble state at acidic/alkaline pHs or a crystalline state at physiological pH. Ueno [6] also reported that CipA shares considerable structural similarity with the B subunit of heat-labile enterotoxin type IIB (PDB ID: 1QB5), which is known to form a pentameric complex. Therefore, proteins with a globular oligonucleotide/oligosaccharide-binding fold domain can exhibit variable oligomeric valency depending on their state.

At pH 10.6, CipA and CipB showed markedly different behaviors in solution. While CipB remained as a stable tetramer, CipA formed a much larger complex, such as a decamer around 117 kDa, suggesting that CipA undergoes extensive non-covalent self-associations in basic pH environments. The notable difference in CipA’s solution behavior between acidic (pH 3.5) and basic (pH 10.6) conditions is most likely attributable to pH-dependent alterations in surface charge distribution or the exposure of new interaction sites that facilitate self-association only at alkaline pH. The biological significance of the covalently linked CipA adducts remains undetermined. Considering the proposed nutritional roles for CipA and CipB crystals, it is notable that CipA and CipB crystals dissolve into trimers and tetramers at the gastric pH range, while displaying divergent dissolution kinetics and varying non-covalent and covalent oligomeric states at the alkaline pH of the nematode intestinal lumen.

From an engineering standpoint, CipA’s susceptibility to oxidative adduct formation presents both challenges and opportunities. Uncontrolled oxidation can hinder reproducibility and quality control during downstream processing of CipA-based materials in alkaline settings. Conversely, deliberate crosslinking of designated Cys-Cys pairs could potentially stabilize oligomeric assemblies or modify solubility, contingent upon the precise identification and regulation of participating cysteine pairs and relevant reaction parameters.

### 3.4. The phase separation of CipA and CipB into distinct orthogonal crystals agrees with prevailing homotypic interactions

Another key finding was that, when CipA and CipB were co-expressed in the same mammalian cytosol, they underwent phase separation simultaneously and formed two discrete crystal populations that retained their characteristic morphologies without mixing. This “demixing” behavior indicates that homotypic interactions (CipA–CipA and CipB–CipB) are energetically favored over heterotypic (CipA–CipB) and non-specific interactions. Given the low sequence identity between CipA and CipB, it is plausible that differences in surface properties and packing constraints disfavored the formation of a shared lattice. The CipA/CipB pair therefore represents an orthogonal self-assembly system in which two high-concentration self-assembling proteins occupy the same compartment yet partition into separate phases. A series of co-expression experiments further demonstrated the robustness of CipA and CipB crystallization in the crowded cytosolic environment, even in the presence of other unrelated crystallizing proteins, such as CRYGD-R37S [17] and PAK4-cat (278-591) [24]. Notably, the formation of three independent crystals during the simultaneous co-expression of CipA, CipB, and PAK4-cat (278-591) further illustrated their strong homotypic interactions that defy perturbation by other proteins.

CipA and CipB have attracted attention as scaffolding proteins capable of organizing heterologous proteins into crystalline inclusions in bacterial systems for metabolic engineering and bioactive metabolite synthesis [7–15]. These inclusions can also be isolated from bacteria as bioactive materials to perform biochemical reactions *in vitro*. The present study extends the potential of the Cip-based enzyme scaffolding approach into eukaryotic cellular contexts by revealing that CipA and CipB can generate abundant, discrete, and significantly larger cytosolic crystals in mammalian cells. Considering that mammalian intracellular crystals, such as those formed by PAK4/Inka1, have already been used in structural determination through *in-cellulo* X-ray crystallography [24], enzyme scaffolding via the “crystalline flask” method [24], and solid biomaterial production [26], CipA/CipB expands the selection and scope of proteins available to construct large intracellular crystals with diverse functionalities. Furthermore, the simultaneous phase separation of CipA, CipB, and PAK4-cat into three distinct crystalline phases within the same cytosol broadens the programmability of the CipA/CipB scaffold system for synthetic biology applications. If combined with the PAK4-cat system, CipA/CipB scaffolds could potentially facilitate the creation of three concurrent crystalline compartments for storage or reaction, where each crystal serves as a platform for different enzymes in the shared cytosolic space while maintaining spatial segregation. This capability should enable precise separation of enzyme activities or cargos within a single subcellular compartment, allowing better control of metabolic pathways and multi-step reactions in engineered mammalian cells. To advance these findings within mammalian synthetic biology, further work is needed to validate whether these crystals can effectively incorporate heterologous enzymes as fusion partners in the cytosol of mammalian cells while preserving crystal integrity and enzymatic activity.

## 4. Materials and Methods

### 4.1. Detection antibodies and reagents

Mouse anti-NUP153 (clone QE5) was procured from Abcam (cat. Ab24700). Rabbit anti-giantin was obtained from Covance (cat. PRB-114P). Unless otherwise noted, all chemicals and biological reagents were sourced from Sigma-Aldrich.

### 4.2. Expression constructs

The amino acid sequences of *Photorhabdus luminescens* CipA and CipB are described in UniProt KB under entries Q56816 and P96969, respectively. Similarly, the corresponding sequences for CipA and CipB of *Photorhabdus laumondii* can be found under UniProt KB IDs Q7N6H4 and Q7N9Z4. For optimal recombinant gene expression in human cells, nucleotide sequences were reverse translated from these amino acid templates using a publicly accessible codon optimization tool provided by GENEWIZ (South Plainfield, NJ, USA). The construct encoding human γ-crystallin D with the R37S point mutation (CRYGD-R37S) has been previously characterized [17], and the sequence of PAK4-cat (residues 278-591) was reported in an earlier publication [24]. All recombinant DNA constructs underwent sequence verification and were then seamlessly subcloned into Golden Gate-compatible pTT^®^5 mammalian expression vectors licensed from the National Research Council of Canada.

### 4.3. Cell culture and transfection

HEK293-6E cell line (HEK293) was obtained from the National Research Council of Canada. Suspension cultures of HEK293 cells were maintained at 37 °C with 5% CO_2_ in a humidified incubator, using FreeStyle™ 293 Expression Medium (Thermo Fisher Scientific). Cultures were grown in vented cap Corning^®^ Erlenmeyer flasks on shaker platforms operated at 130-135 rpm. Transfection of the expression construct into HEK293 cells was performed using the well-established polyethyleneimine transfection method. At 24 hours post-transfection, Difco yeastolate cell culture supplement (BD Biosciences) was added to the suspension cell cultures. For co-transfection experiments involving two genes, 1:1 plasmid DNA ratio (50% dosage per gene) was used without altering the total DNA amount. Similarly, for three-gene co-transfections, a 1:1:1 plasmid DNA ratio (33% dosage per gene) was employed.

### 4.4. Microscopy

Suspension-cultured transfected cells were seeded onto poly-D-lysine-coated glass coverslips (Neuvitro Corporation) 48 hours after transfection. These cells were then kept in static culture at 37 °C up to day-6 post-transfection. For certain experiments, transfected cells were seeded onto glass coverslips 24 hours after transfection and moved to a CO_2_ incubator with a cooling capability (ESCO CelCulture®) maintained at 27 °C. At specific time points following transfection, the cells were fixed with 4% paraformaldehyde in 100 mM sodium phosphate buffer (pH 7.2) for 30 min at room temperature. Fixed cells were either directly examined using differential interference contrast (DIC) and phase contrast microscopy or prepared for indirect immunofluorescent microscopy. For immunostaining, cells were permeabilized for 15 minutes in phosphate-buffered saline containing 0.4% saponin, 1% bovine serum albumin, and 5% fish gelatin, then incubated with primary antibodies for 60 minutes. After three washes in the permeabilization buffer, secondary antibody labeling was carried out for another 60 minutes. Stained coverslips were mounted onto slides using Vectashield mounting medium (Vector Laboratories) and analyzed with a Nikon Eclipse 80i microscope equipped with a 60× or 100× CFI Plan Apochromat oil objective lens and Chroma FITC-HYQ or Texas Red-HYQ filters. DIC and fluorescent images were acquired via an ORCA-Fusion digital CMOS camera (Hamamatsu) and Nikon Elements BR software. Phase contrast microscopy was performed with a Zeiss AXIO Imager.A1 fitted with an AxioCam CCD camera.

### 4.5. Cell lysis and SDS-PAGE analysis

On day-3 post-transfection, an aliquot of suspension cell culture was removed from shake flasks, and cell pellets were harvested via centrifugation at 1,000 g for 5 minutes. Cell pellets were then lysed directly in 1× sodium dodecyl sulfate (SDS) sample buffer (Thermo Fisher Scientific) containing 5% (v/v) β-mercaptoethanol and heat-treated at 75 °C for 5 minutes. Sample loading was normalized by analyzing whole cell lysates equivalent to 12,000–12,500 cells per lane. SDS-PAGE was performed using a NuPAGE 4–12% Bis-Tris gradient gel and MES SDS buffer system (both from Thermo Fisher Scientific). The gels were then stained with InstantBlue® Coomassie Protein Stain (Abcam).

### 4.6. Isolation and solubilization of intracellular protein crystals

To isolate protein crystals from transiently transfected HEK293 cells, cells were collected by low-speed centrifugation (1,500 g, 7 min) 72 hours after transfection. Intracellular crystals were released by lysing the cell pellets in Hepes-buffered saline (HBS) (20 mM Hepes, 150 mM KCl, 2 mM EDTA, pH 7.4) containing 1% (v/v) Triton X-100, followed by incubation on ice for 15 minutes. The released crystals were collected by centrifuging at 1,500 g for 7 min. Crystal pellets were resuspended in HBS (pH 7.4) containing 1% (v/v) NP-40 by gentle pipetting, incubated on ice for 15 minutes, and centrifuged for 7 min at 1,500 g. The collected crystal pellets were washed using an excess volume of HBS (pH 7.4) with gentle pipetting. After the final wash, crystals were resuspended in HBS (pH 7.4) and were visually inspected under the microscope.

To evaluate the solubility of protein crystals, the slurry containing CipA or CipB crystals in HBS (pH 7.4) was divided into three equal aliquots and spun at 3,000 g for 10 minutes. After the supernatant was removed, each crystal pellet was resuspended in one of the three different buffers: (1) HBS at pH 7.4; (2) 100 mM sodium bicarbonate buffer at pH 10.6; (3) 0.5% acetic acid, 150 mM NaCl at pH 3.5, and incubated at 4°C for up to 19 days. The purity and amount of solubilized CipA and CipB were analyzed by SDS-PAGE under either reducing or non-reducing conditions. Rough protein crystal yields were estimated based on starting cell numbers and the final protein concentration and volume of solubilized protein.

### 4.7. Analytical Size Exclusion Chromatography (SEC)

The quality and purity of solubilized CipA and CipB, derived from isolated intracellular crystals, were evaluated using analytical SEC on an Agilent 1200 series HPLC equipped with a Superdex 200 column (GE Healthcare). Analyses were performed either in a pH 3.5 buffer (0.5% acetic acid, 150 mM NaCl) or a pH 10.6 buffer (100 mM sodium bicarbonate), both running at a flow rate of 0.45 ml/min for an 8-minute run time. Protein samples were clarified by centrifugation at 16,000 g for 5 min before injecting 10 μl of each sample at ∼5 mg/ml. The relative molecular weights were compared to a Gel Filtration Standard (cat. 1511901) from Bio-Rad.

### 4.8. Size-exclusion chromatography coupled to multi-angle light scattering (SEC–MALS)

SEC–MALS was performed using an Agilent 1290 UPLC system interfaced with a Wyatt DAWN MALS detector and an Optilab refractive index detector. Chromatographic control and data acquisition for the LC system were carried out using Chromeleon software, while MALS data were collected and analyzed using ASTRA 8 (Wyatt Technology). The top 20% of each peak was used for molecular weight calculations. The mobile phase consisted of either 0.5% acetic acid, 150 mM NaCl, pH 3.5 or 100 mM bicarbonate, pH 10.6. Samples were separated on a Superdex 200 Increase 5/150 column at a flow rate of 0.3 ml/min with a total run time of 25 min. Prior to analysis, samples were centrifuged at 14,000 rpm for 5 min to remove insoluble aggregates. A 40 µl aliquot of each sample was injected for analysis.

## 6. Acknowledgements

The authors thank Amgen’s Discovery Protein Science molecular cloning team (Mina Mostafavi and Saee Nasikkar) for gene fragment ordering, assembly, sequencing, and plasmid preparation. HH is personally grateful to Yoko Azumi for constant support.

## 8. Data Availability Statement

The authors confirm that the data supporting the findings of this study are available within the article and its supplementary material.

## 9. CRediT authorship contribution statement

**Haruki Hasegawa:** Conceptualization, Resources, Data curation, Investigation, Methodology, Visualization, Project administration, Writing – original draft, Writing – review & editing **Songyu Wang:** Resources, Data curation, Visualization **Emma Pelegri-O’Day:** Resources, Data curation, Visualization

## 11. Figure legends

**Supplement 1.**
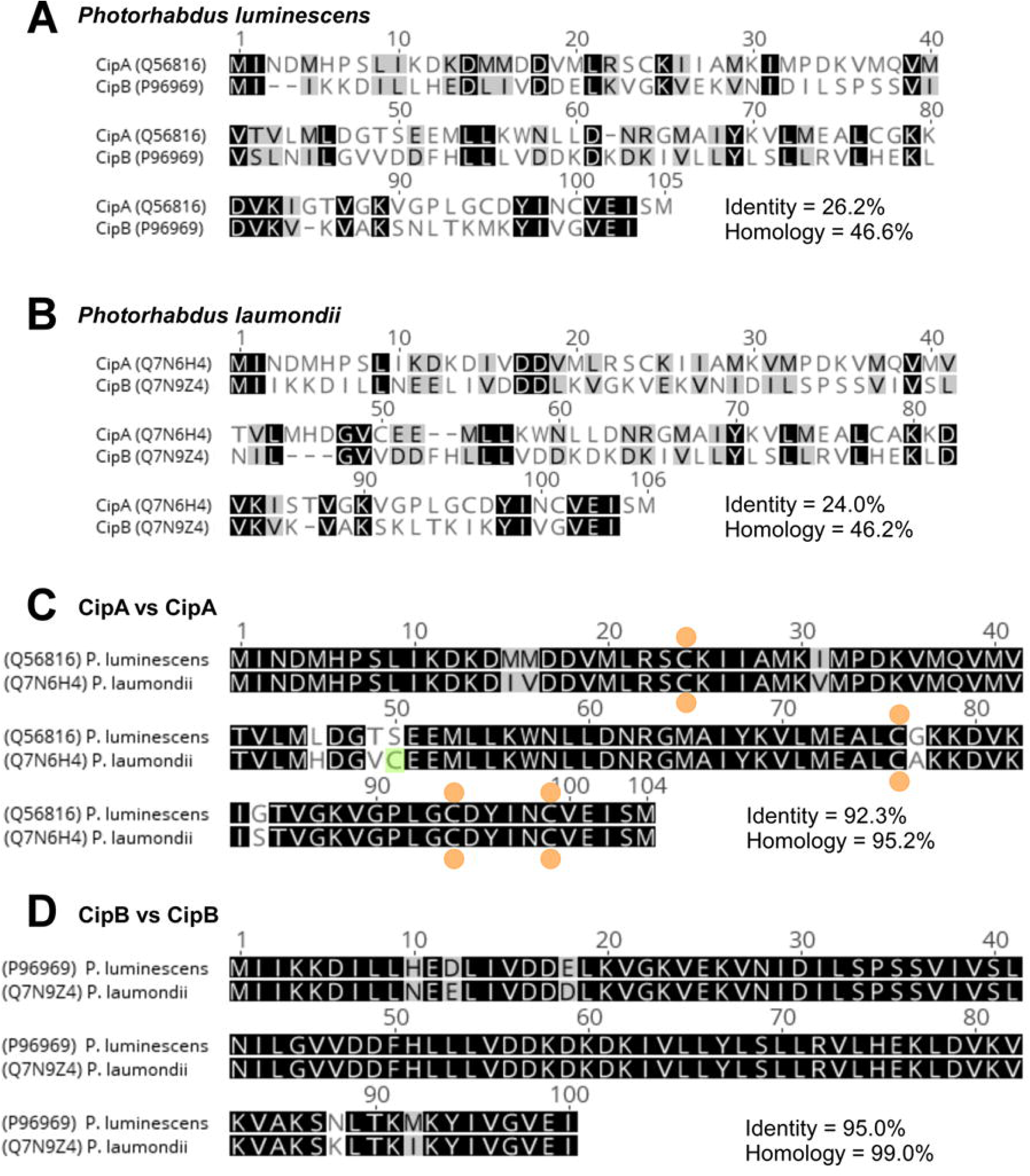
Amino acid sequence alignments of CipA and CipB. (A) Alignment of *P. luminescens* CipA (UniProt Q56816) and CipB (UniProt P96969) amino acid sequences. Identical amino acids and similar amino acids are shaded in black and grey background, respectively. (B) Alignment of *P. laumondii* CipA (UniProt Q7N6H4) and CipB (UniProt Q7N9Z4) amino acid sequences. (C) CipA sequences from *P. luminescens* and *P. laumondii* are compared. Conserved four cysteine residues are marked by orange circles. Non-conserved Cys-50 present only in *P. laumondii* CipA is shaded in green. (D) Likewise, the alignment of CipB sequences from *P. luminescens* and *P. laumondii*. CipB proteins are devoid of cysteine residue.

## References

1. Bintrim, S.B. and J.C. Ensign, Insertional inactivation of genes encoding the crystalline inclusion proteins of Photorhabdus luminescens results in mutants with pleiotropic phenotypes. J Bacteriol, 1998. 180(5): p. 1261–9.

2. Forst, S., et al., Xenorhabdus and Photorhabdus spp.: bugs that kill bugs. Annual review of microbiology, 1997. 51(1): p. 47–72.

3. Ciche, T.A., et al., A Phosphopantetheinyl transferase homolog is essential for Photorhabdus luminescens to support growth and reproduction of the entomopathogenic nematode Heterorhabditis bacteriophora. J Bacteriol, 2001. 183(10): p. 3117–26.

4. Bowen, D.J. and J.C. Ensign, Isolation and characterization of intracellular protein inclusions produced by the entomopathogenic bacterium Photorhabdus luminescens. Appl Environ Microbiol, 2001. 67(10): p. 4834–41.

5. You, J., et al., Nutritive significance of crystalline inclusion proteins of Photorhabdus luminescens in Steinernema nematodes. FEMS Microbiol Ecol, 2006. 55(2): p. 178–85.

6. Abe, S., et al., Cell-free protein crystallization for nanocrystal structure determination. Sci Rep, 2022. 12(1): p. 16031.

7. Wang, Y., R. Heermann, and K. Jung, CipA and CipB as Scaffolds To Organize Proteins into Crystalline Inclusions. ACS Synth Biol, 2017. 6(5): p. 826–836.

8. Huo, Y.X., et al., CipA-mediating enzyme self-assembly to enhance the biosynthesis of pyrogallol in Escherichia coli. Appl Microbiol Biotechnol, 2018. 102(23): p. 10005–10015.

9. Zhao, L., et al., In Vitro Biosynthesis of Isobutyraldehyde Through the Establishment of a One-Step Self-Assembly-Based Immobilization Strategy. J Agric Food Chem, 2021. 69(48): p. 14609–14619.

10. Park, S.Y., et al., Metabolic engineering of Escherichia coli with electron channelling for the production of natural products. Nature Catalysis, 2022. 5(8): p. 726–737.

11. Wang, P., et al., Construction of a Protein Crystalline Inclusion-Based Enzyme Immobilization System for Biosynthesis of PAPS from ATP and Sulfate. ACS Synth Biol, 2023. 12(5): p. 1487–1496.

12. Li, J., et al., Construction of immobilized enzyme cascades for the biosynthesis of nucleotide sugars UDP-N-acetylglucosamine and UDP-glucuronic acid. Systems Microbiology and Biomanufacturing, 2024. 4(3): p. 895–905.

13. Wang, Y.W., et al., Carrier-free immobilized enzymatic reactor based on CipA-fused carbonyl reductase for efficient synthesis of chiral alcohol with cofactor self-sufficiency. Int J Biol Macromol, 2024. 276(Pt 1): p. 133873.

14. Yang, F., et al., Development of Escherichia coli by combining channel engineering and energy engineering for the production of L-histidine. Chemical Engineering Journal, 2025. 520: p. 165521.

15. Zhang, Y., et al., Biocatalytic Reductive Amination for In Vitro Biosynthesis of the Amaryllidaceae Alkaloid Precursor. ChemCatChem, 2025. 17(6): p. e202401811.

16. Wei, X., W. Zhang, and B. Zhu, Harnessing a Genetically Engineered Self-Assembling Protein Biosorbent for Efficient and Selective Rare Earth Element Recovery. Environ Sci Technol, 2025. 59(34): p. 18213–18224.

17. Hasegawa, H., Simultaneous induction of distinct protein phase separation events in multiple subcellular compartments of a single cell. Exp Cell Res, 2019. 379(1): p. 92–109.

18. Schonherr, R., J.M. Rudolph, and L. Redecke, Protein crystallization in living cells. Biol Chem, 2018. 399(7): p. 751–772.

19. Mudogo, C.N., et al., Protein phase separation and determinants of in cell crystallization. Traffic, 2020. 21(2): p. 220–230.

20. Hasegawa, H., et al., Light chain subunit of a poorly soluble human IgG2lambda crystallizes in physiological pH environment both in cellulo and in vitro. Biochim Biophys Acta Mol Cell Res, 2021. 1868(9): p. 119078.

21. Hasegawa, H., Demixing of four simultaneously co-expressed phase-separating proteins in the endoplasmic reticulum lumen. Biosci Rep, 2025. 45(9): p. 481–90.

22. Machado, R.A.R., et al., Whole-genome-based revisit of Photorhabdus phylogeny: proposal for the elevation of most Photorhabdus subspecies to the species level and description of one novel species Photorhabdus bodei sp. nov., and one novel subspecies Photorhabdus laumondii subsp. clarkei subsp. nov. Int J Syst Evol Microbiol, 2018. 68(8): p. 2664–2681.

23. Hasegawa, H., Temperature-dependent intracellular crystallization of firefly luciferase in mammalian cells is suppressed by D-luciferin and stabilizing inhibitors. Exp Cell Res, 2024. 440(1): p. 114131.

24. Baskaran, Y., et al., An in cellulo-derived structure of PAK4 in complex with its inhibitor Inka1. Nat Commun, 2015. 6: p. 8681.

25. Hasegawa, H., et al., In vivo crystallization of human IgG in the endoplasmic reticulum of engineered Chinese hamster ovary (CHO) cells. J Biol Chem, 2011. 286(22): p. 19917–31.

26. Li, T.L., et al., Engineering a Genetically Encoded Magnetic Protein Crystal. Nano Lett, 2019. 19(10): p. 6955–6963.

